# The Electrophysiological Markers of Statistically Learned Attentional Enhancement: Evidence for a Saliency Based Mechanism

**DOI:** 10.1101/2023.03.14.532560

**Authors:** Dock H. Duncan, Dirk van Moorselaar, Jan Theeuwes

## Abstract

It has been well established that attention can be sharpened through the process of statistical learning - whereby visual search is optimally adapted to the spatial probabilities of a target in visual fields. Specifically, attentional processing becomes more efficient when targets appear at high relatively to low probability locations. Statistically learned attentional enhancement has been shown to differ behaviorally from the more well studied top-down and bottom-up forms of attention; and while the electrophysiological characteristics of top-down and bottom-up attention have been well explored, relatively little work has been done to characterize the electrophysiological correlates of statistically learned attentional enhancement. In the current study, EEG data was collected while participants performed the additional singleton task with an unbalanced target distribution. Encephalographic data was then analyzed for two well-known correlates of attentional processing – alpha lateralization and the N2pc component. Our results showed that statistically learned attentional enhancement is not characterized by alpha lateralization, thereby differentiating it from top-down enhancement. Yet targets at high probability locations did reliably produce larger N2pc amplitudes, a known marker of increased bottom-up capture due to higher target-distractor contrasts. These results support an interpretation of the probability cuing effects where the improved processing of targets at expected locations is mediated by a saliency-based mechanism – boosting the salience of targets appearing at high-probability locations relative to those at low-probability locations.

**Significance statement:** Things are easier to find when you have a good idea of where they should be – e.g. shoes on the floor and birds in the sky. Expectations of where things are likely to be found can be implicitly learned without much, if any, awareness. Until now, little was known about how these implicit spatial biases change the representation of items in the brain. In the current work, we present EEG recordings which suggest that the brain may represent items in common locations as more salient than in other locations in space. These findings inform how the brain represents implicit search expectations; supporting a model where items in expected areas in space capture attention more frequently because they are represented by the brain as more salient.

## Intro

When we look upon a scene, several factors, - including low level feature contrasts within a scene (a red stop sign automatically pops out from a grey background) and our current search priorities (we notice street names when trying to find a specific address) - coalesce to determine how attention is deployed. Traditionally it was assumed that attentional selection was solely determined by the interaction between these bottom-up (salience-driven) and top-down (goal-directed) factors (Koch & Ullman, 1984; Wolfe, 1994; Egeth & Yantis, 1997; Theeuwes, 2010). However, it is becoming increasingly appreciated that attentional deployment is also heavily influenced by past experiences with the current stimuli and search environment (Awh et al., 2012; Geng & Behrmann, 2005; Jiang, 2018; Theeuwes, 2019); think for instance how driving down a familiar road is a qualitatively different experience than driving down an unfamiliar one. The influence of past experiences on future attentional behavior is known as selection history; a category of phenomena including intertrial priming, reward learning, and statistical learning (Anderson et al., 2021; Failing & Theeuwes, 2018; Theeuwes, 2018). Recent studies have demonstrated that through statistical learning in particular, attentional priority settings are optimally adjusted to regularities in the environment; and counter to top-down goals induced by explicit instructions, this learning may occur without intention and often without observers being aware (for a review, see Theeuwes et al., 2022).

It has been known for some time that when targets appear more often in one location than in other locations, participants learn this structure and detect targets faster at this high probability location than at low probability locations (Geng & Behrmann, 2002, 2005). This attentional effect has been shown to scale monotonically with the probability that targets appear in one particular spatial location (Zhang et al., 2022), demonstrating that through statistical learning the weights within an assumed visual spatial priority map are adjusted optimally, resulting in efficient selection priorities (Theeuwes et al., 2022).

While top-down and bottom-up and influences on attentional selection have been intensively studied both at a behavioral (Luck et al., 2021) and neural level (Desimone & Duncan, 1995; Fecteau & Munoz, 2006; Itti & Koch, 2001), surprisingly little neuroimaging work has been done to elucidate the cognitive mechanisms underlying statistically learned attentional enhancement. It thus remains unclear to what extent the well-known speedup effect of probability cuing is the result of an implicit proactive shift of covert attention, similar to anticipatory covert shifts of attention induced by top-down goals, or alternatively reflects mechanisms that are better characterized as a form of bottom-up selection.

If learning about probable target locations is implemented via similar mechanisms that drive top-down attention, it should be reflected in oscillatory neuronal activity within the alpha-band, which decreases over contralateral occipital cortex relative to the attended location in response to a central attention cue (Rihs et al., 2007; Sauseng et al., 2005; Worden et al., 2000). Alternatively, if probability cuing does not result in a proactive shift of covert attention, but instead changes neural processing of the search display, this should be indexed via amplitude modulations in the N2pc, a contralateral negativity measured with EEG which indexes visual selection and has shown to increase as relative target saliency increases (Mazza et al., 2009; Töllner et al., 2011). Such a finding would concur with recently proposed models of statistical learning whereby attentional enhancement is mediated by changing weights on the attentional priority map, modifying latent neural excitability in the high-probability locations in space (Duncan et al., 2022; Ferrante et al., 2023; van Moorselaar et al., 2020). To dissociate between these alternatives, here we recorded scalp level EEG while participants performed a modified version of the additional singleton task with unbalanced target distributions. This allowed us to characterize the electrophysiological correlates of statistically learned attentional enhancement, and to compare this neural signature to those known to characterize top-down and bottom-up attention.

## Methods

### Participants

This article reports novel analyses of data previously presented in Duncan, van Moorselaar and Theeuwes (2022). The sample size of 24 participants was thus selected based on the expected effect size from that project; however, this sample size also exceeds that of what is typically used in other ERP and time-frequency studies (Foster et al., 2020; Wang et al., 2019). Out of the original sample, two participants were excluded during reanalysis because their eye tracking data revealed that they deviated from fixation on more than 30% of trials (see eye tracking methods below – note this was not an issue for published results as they focused on a different window of interest). The final dataset of analysis thus consisted of 22 participants (17 female, average age 21). The study was conducted in the Brain and Behavior labs at Vrije Universiteit Amsterdam; all participants indicated normal or corrected-to-normal vision and reported no history of cognitive impairments. All participants additionally provided written informed consent prior to participation and were compensated with 25 euros or equivalent course credit. The study conformed to the Declaration of Helsinki and was approved by the Ethical Review Committee of the Faculty.

### Apparatus and Stimuli

Participants were seated in a sound-attenuated room with dim lighting 60cm away from a 23.8 inch, 1920×1080 pixel ASUS ROG STRIX XG248 LED monitor with a 240hz refresh rate, which displayed all stimuli. The behavioral task was programmed using OpenSesame (Mathot et. al., 2012) and incorporated functions from the Psychopy library of psychophysical tools (Pierce, 2007). EEG data were recorded using a 64-electrode cap with electrodes placed according to the international 10–10 system (Biosemi ActiveTwo system; biosemi.com), with two earlobe electrodes used as offline reference, and default Biosemi settings at a sampling rate of 512 Hz. External electrodes placed ~2 cm above and below the right eye, and ~1 cm lateral to the left and right lateral canthus were used to record vertical (VEOG) and horizontal EOG (HEOG). Eye-tracking data were collected using an Eyelink 1000 (SR research) eye tracker that tracked both eyes. To maintain stability, participants used a headrest positioned 60cm from the screen. Eye sampling varied between participants at 500, 1000, and 2000hz due to the different EEG labs used having different versions of the Eyelink 1000 with varying sampling limits. All participant data was later aligned with the EEG data during preprocessing. Calibration occurred before the first block and at the halfway point for all participants. Additional calibrations were performed for some participants due to subtle changes in resting position in the chinrest or if noticeable drift in the eye signal was suspected by the experimenter.

All stimuli were presented on a black background. The fixation point (~1.1°) was that shown by Thaler et al. (2013), to enforce stable fixation. The search display consisted of eight equally spaced shapes presented on an imaginary circle centered at fixation (radius = ~4.8°). Shapes could either be diamonds (82×82 px or ~ 2.1°×2.1° square rotated 45 degrees) or circles (diameter 90px, ~2.4°) and could be colored either red (rgb 255,0,1) or green (rgb 0,128,0). All shapes also contained a horizontally or vertically oriented white line (70px; ~1.8°, rgb 255,255,255). The displays were rendered such that each display contained one unique shape (i.e., the target), and on a subset of trials one of the homogeneous shapes had a unique color (i.e., singleton distractor).

### Procedure and Design

The task was a combination of the additional singleton task (Theeuwes, 1992) and the probability cuing task (Geng & Behrmann, 2005) where targets were presented more frequently in certain regions in space (high-probability locations, see Figure 1A for an example of a task display). This sort of task has been shown to lead to an attentional facilitation at the high-probability location relative to the other low-probability locations (Ferrante et al., 2018; Gao & Theeuwes, 2020a; Geng & Behrmann, 2005; Huang et al., 2022). Trials began with a fixation point onset, which lasted between 1.3 and 1.7 seconds. On 50% of trials, a high contrast, task-irrelevant ping was presented for 200ms sometime between 700-900ms after fixation onset (see Duncan et al., 2022 for a neural analysis centered on these perturbations). These trials were first analyzed separately from trials in which the screen remained blank for the entire intertrial interval in our analysis of intertrial alpha. After that, the search display appeared and participants were instructed to find the unique shape singleton target and indicate the orientation of the line within by pressing either the ‘left’ direction key (for ‘horizontal’) or the ‘up’ direction key (for ‘vertical’). The search arrays remained on screen for 2.5 seconds, or until participants provided a response. On a subset of trials (70%) the search display also contained a singleton distractor, which had to be ignored.

**Figure 1:**
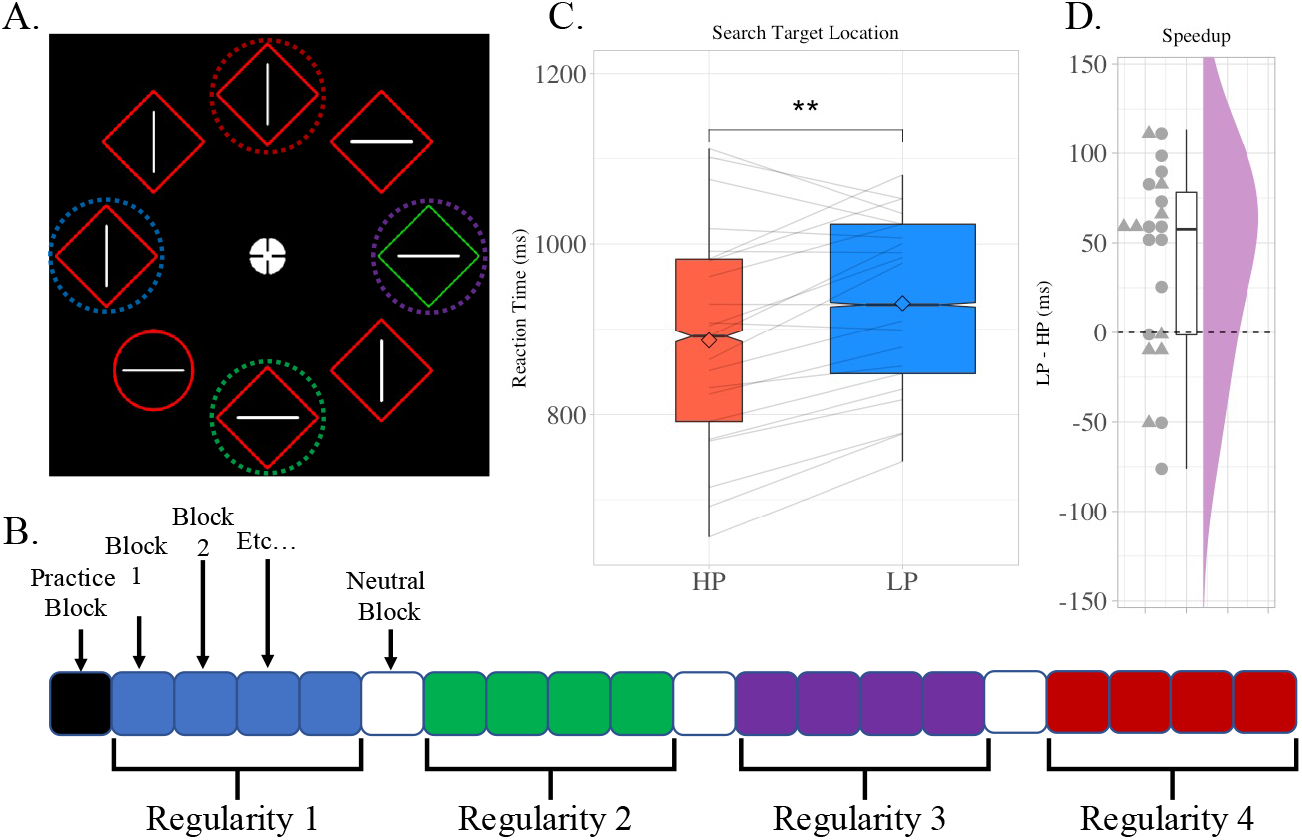
A) An example of the additional singleton search display. In this task, participants look for a unique shape and report the orientation of the line held within this shape singleton. While they perform this task, a color singleton is occasionally presented as a distractor. In the current study, an imbalance in the target location distribution evoked a probability cuing effect, where participants were implicitly trained to expect targets to appear in certain locations in space – as reflected by faster reaction times. Denoted by the dotted circles are the four possible high-probability locations in space. Note that in the actual experiment, the four dotted circles did not appear. The colors of these circles correspond to the hypothetical block order shown in: B) Visual representation of the experiment’s block structure. During the experiment, the high-probability target location occasionally shifted such that each participant would encounter a series of blocks where each of the four possible high-probability locations was active. These locations were the four cardinal locations (top, bottom, left and right). High-probability locations would be stable for four blocks at a time before switching (symbolized by the four squares of the same color). Note that the four high-probability locations were counterbalanced between the participants such that no two participants encountered the four locations in the same order (the four locations shown in A are merely an example of one possible presentation order). In between high-probability blocks, three neutral blocks (represented by the white squares) were inserted. These blocks were meant to aid in the extinction of the previous probability cuing effect. C) Reaction times when targets were presented in high-probability (HP, red) trials or low-probability (LP, blue) trials. Participants were reliably faster on HP trials than LP trials. D) The individual speedup effect for each participant – calculated by taking each participants average reaction time on LP trials and subtracting their average reaction time for HP trials. Triangles represent participant averages for participants that indicated some awareness of the target regularity. Circles are participant averages for participants that reported no awareness of the target regularity.

Critically, targets did not appear at random locations across trials, but instead followed an undisclosed distributional regularity where they were disproportionately more likely to appear in one location than any other. These high-probability (HP) target locations could either be the left, right, top or bottom most positions in the search array. Furthermore, these locations shifted periodically (i.e., high-probability locations were used for four consecutive blocks before switching) during the experiment such that one location was only the high probability location for a certain amount of time in each experiment session (see Figure 1B; order counterbalanced across participants^1^). Thus, counter to typical probability cuing studies, condition averages reflected data across multiple display locations, rather than a single location benefit. Neutral blocks, in which targets appeared equally frequently at all 8 locations, were inserted in between switches to give participants an opportunity to un-learn the previous high-probability location (Britton & Anderson, 2020; Duncan & Theeuwes, 2020; Valsecchi & Turatto, 2021) before moving on to a new high-probability condition.

The experiment itself consisted of 19 blocks of 56 trials each, plus one full practice block. In between blocks participants received feedback on their performance (i.e., mean reaction time (RT) and accuracy), and were encouraged to take a short break. Upon completing the experiment, participants were asked whether they noticed any regularity to the target presentation. They were next asked to indicate where they believed the target was most likely to appear in the final block of the experiment, and finally they were asked if there were any other locations that they believed held the target more frequently at any time in the experiment.

### EEG acquisition and Preprocessing

After re-referencing all EEG data to the average of the two earlobe electrodes, a Hamming windowed FIR filter was used to high-pass filter the data at 0.1 Hz to remove slow signal drifts. The continuous EEG signal was then segmented into epochs from 1500ms before search onset to 1000ms after search display onset. A different time window of −1000ms – 500ms was used to reject trials. Malfunctioning electrodes marked as bad by the experimenter during recording were temporarily removed before subsequent trial rejection and artefact correction. To identify EMG contaminated epochs, an adapted version of an automated trial-rejection procedure was used, as implemented in Fieldtrip (Oostenveld et al., 2011). Muscle activity was specifically captured using a 110-140 Hz band-pass filter, and variable z-score thresholds per subject were allowed based on within-subject variance of z-scores (Vries et al., 2017). Rather than immediately removing epochs exceeding the z-score threshold, the five electrodes that contributed most to accumulated z-score within the time period containing the marked EMG artefact were first identified. An iterative procedure was then used to interpolate the five worst electrodes per marked epoch using spherical splines (Perrin et al., 1989), checking whether that epoch still exceeded the determined z-score threshold after interpolation. Epochs were only dropped when after interpolation of the five worst electrodes the z-score threshold was still exceeded. Independent Component Analysis (ICA) was then applied to the epoched data that had been high-pass filtered at 1 Hz using the ‘picard’ method as implemented in MNE, to remove eye-blink components from the cleaned data. Finally, identified malfunctioning electrodes were interpolated using spherical splines.

### Eye-tracking exclusions

During the experiment, participants were instructed to keep their eyes locked on fixation at all times. To ensure this was the case, eye tracking data was recorded, with real time feedback given to participants when their eyes deviated from fixation in the form of a high-pitched beep. This eye tracking data was further used offline to exclude trials in which an eye movement was made. Using custom scripts, x, y positions were linearly interpolated in between the start and end of an eyeblink (±20ms). Trials with a fixation deviation >1° of visual angle in a segment of continuous data of at least 40ms in specific time windows (−1000 – 0ms and −200 – 350ms for the time-frequency and ERP analysis respectively). In case, a trial had missing eye tracker data, the HEOG recorded signal was examined instead for sudden jumps in the recorded signal – a known marker of saccades. A step method was used, with a window of 200ms, a threshold of 15μV and a step size of 10ms. In total, ~7% of trials in the ERP analysis and ~8%of trials in the alpha analysis were excluded due to eye movements (and as noted above two participants from the original sample were excluded as more than 30% of trials were marked for exclusion).

### Behavioral Analysis

Because only the left and right high-probability conditions were considered in our EEG analyses, analysis of targets at high probability locations was restricted to these locations. Low-probability trials, on the other hand, could be drawn from any block, regardless of where the current high-probability location was. Trials were excluded for analysis if the participant provided an incorrect response, of if response time on that trial was more than 2.5 standard deviations away from the participant mean reaction time.

### TF analysis

For our TF analysis, separate analyses were conducted on trials in which a salient white ping was presented in the intertrial period, as well as on trials in which the screen remained static (trials were 50/50 across these conditions, see procedure and design). Following these separate analyses, all trials were collapsed into a single analysis. Our analyses were further restricted only to blocks in which the high-probability target was on the horizontal midline (left or right) as alpha lateralization has most commonly been studied in the context of horizontal cues (Klimesch, 2012).

Frequency specific activity in the EEG signal was isolated using Morlet wavelet convolution to decompose the combined EEG signal into 25 logarithmically spaced steps of a frequency range from 4 to 40hz. To create complex Morlet wavelets, for each frequency a sine wave (*e^i2ρft^*, where *i* is the complex operator, *f* is frequency, and *t* is time) was multiplied by a Gaussian (*e^-t^2/2s^2^*, where *s* is the width of the Gaussian). To keep a good trade-off between temporal and frequency precision the Gaussian width was set as:

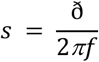

where ð represents the number of wavelet cycles, in 25 logarithmically spaced steps between 3 and 12. Frequency-domain convolution was then applied by multiplying the Fast Fourier Transform (FFT) of the EEG data and Morlet wavelets. The resulting signal was converted back to the time domain using the inverse FFT. Time-specific frequency power, which was down sampled by a factor of 4, was defined as the squared magnitude of the complex signal resulting from the convolution.

To investigate whether the presence of a high-probability location on the left or right of the horizontal midline resulted in a lateralization of alpha-power, we first calculated a lateralization index over the broadband frequency range; this was done by taking frequency-band data for all frequency bins calculated over in contralateral sensors across all time points and subtracting them from those calculated over ipsilateral sensors. This matrix was then divided by the value of adding both ipsilateral and contralateral frequency values together (van Moorselaar et al., 2020):

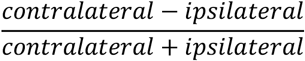

A positive number then would indicate that contralateral power is larger than ipsilateral power, and vice versa for negative numbers. Critically, this index does not require a baseline. Statistical analyses were limited to electrode pairs PO7/8, PO3/4 and O1/2, which were selected on the basis of visual inspection of the topographic distribution of averaged alpha power (8–12 Hz) across the anticipatory time window (−1000 to 0ms) and also were matched to those used in Wang et al. (2019) who previously found alpha lateralization following statistical distractor learning. This analysis was further repeated using only the average alpha band frequencies in both the total alpha band (8-12Hz) as well as separately for upper- and lower-alpha bands (10-13Hz & 7-10Hz respectively).

### ERP analysis

We focused the analysis of target-elicited ERP waveforms on the same electrode pairs PO7/8, PO3/4 and O1/2 that were used in the time-frequency analysis, and that are also often used to study the N2pc (Luck, 2014). Epochs were 30 Hz low-pass filtered and then baseline corrected using a −200 – 0-ms pre-stimulus baseline period. To enable isolation of lateralized target-specific components, the analyses focused on trials in which the target was presented to the left or right of fixation, while the distractor was either absent or present on the vertical meridian – thereby evoking no lateralized component (Hickey et al., 2009; Woodman & Luck, 2003). Waveforms evoked by the various search displays were collapsed across left and right visual hemifield and left and right electrodes to produce separate waveforms for contralateral and ipsilateral scalp regions. Lateralized difference waveforms were then computed by subtracting the ipsilateral waveform from the corresponding contralateral waveform. Critically, low probability targets were selected from all trials where the high probability location did not match the target location on that specific trial. Neutral trials were not included in the ERP analysis as they represented the trials in which spatial priority was being unlearned and thus there was no clear a priori expectation on their evoked neural activity. Incorrect trials were also excluded, matching our behavioral analysis. Overall, a roughly matched number of trials were used for high-probability and low-probability conditions (on average; 71 HP and 51 LP trials per participant were used). For our N2pc analysis, we used the same method as in Wang et al. (2019) to identify the window; first finding the most negative point on the condition subtracted difference waves, and then adding 50ms to either side to find a 100ms window of interest. Additionally, as visual inspection suggested that condition differences were most pronounced early in the time window, we separated the N2pc in an early and a late component (Eimer & Kiss, 2007, 2008; Holmes et al., 2009; Woodman & Luck, 1999), by classifying the period post the negativity peak as the late N2pc, and the period preceding the peak as the early-N2pc (see discussion for further details on this).

### Statistics

Behavioral results were analyzed using simple paired t-tests. TF results were analyzed using a cluster permutation test; a method which corrects for multiple comparisons (Cohen, 2014; Maris & Oostenveld, 2007). We specifically used the MNE function permutation_cluster_1samp_test (Gramfort et al., 2014) on both the broadband power data and the averaged alpha channel data in separate analyses (Figure 2B & C respectively). Clusters were identified as contiguous data points in which a t-statistic exceeded the threshold corresponding to a two-sided p-value of 0.05. A null distribution for the test statistic was found via a Monte Carlo randomization procedure which shuffled condition labels across 1024 iterations. Significant clusters in the unpermuted data were identified as those in which the test statistic was larger than the 95^th^ percentile of our simulated null distribution, thus approximating a t-test with an alpha of 0.05. The N2pc analysis used repeated measure analyses of variance (ANOVA’s) and t-tests. To see whether the ipsilateral response was larger than the contralateral response in our window of interest (the N2pc), we had a clear prediction that an N2pc should be observed and thus a one-tailed t-test was used.

**Figure 2:**
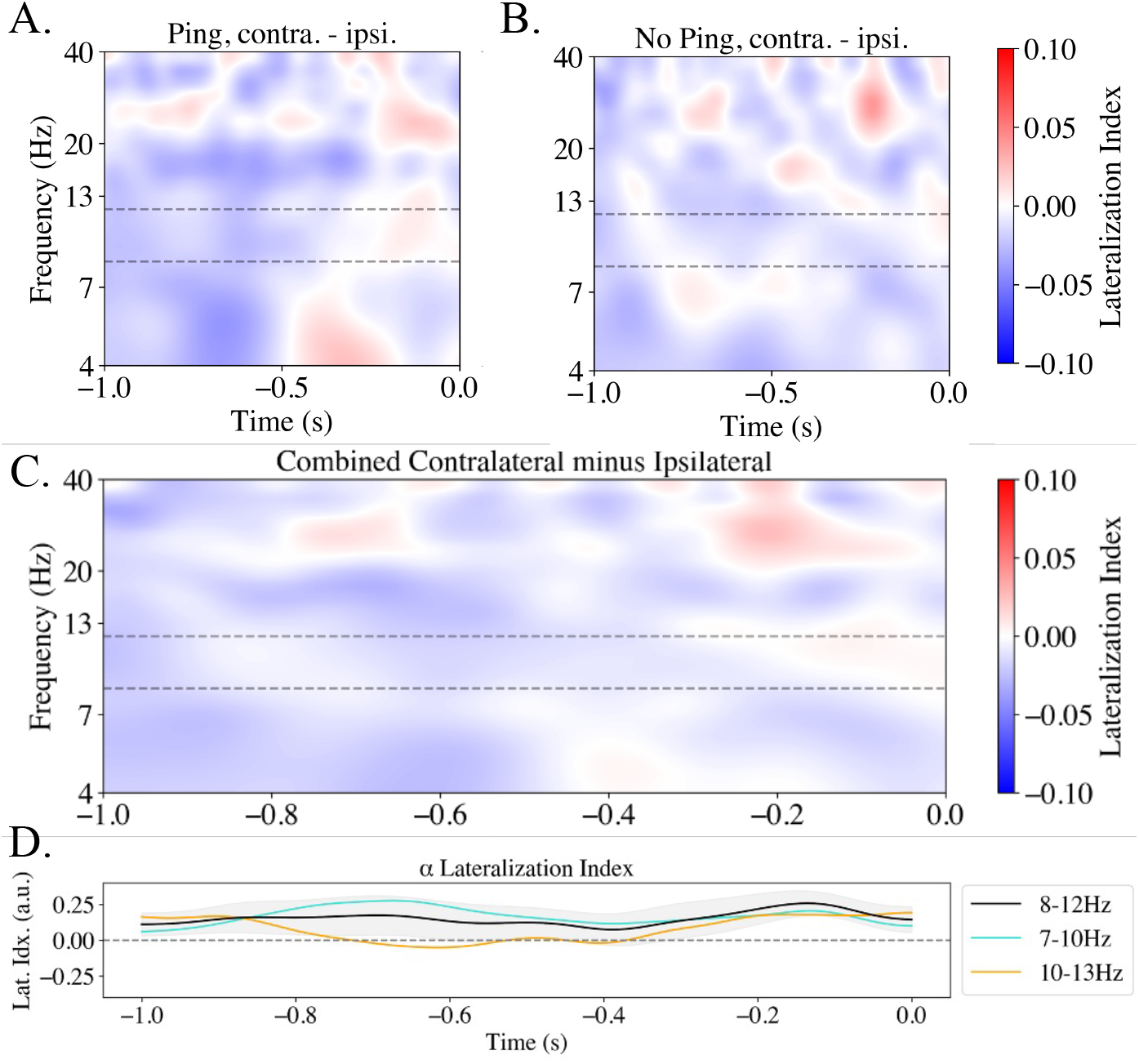
Broadband and alpha power lateralization in the intertrial period from posterior electrodes of interest. Lateralization scores for A) Ping and B) No-ping trials. Lateralization was calculated by subtracting contralateral power from ipsilateral and dividing the result by their combined power. A cluster-based permutation analysis was conducted on these values to identify contiguous regions of significantly positive/negative values. No such clusters were found in either ping nor no-ping trials. Black dotted lines outline region between 7 and 10 Hz used in the main alpha analysis, C) Lateralized power difference between contralateral and ipsilateral electrodes when combining ping and no-ping trials D) Averaged alpha difference for total alpha (black) lower alpha (turquoise) and upper-alpha (orange). Light gray shaded are represents standard error around total alpha condition. A cluster permutation test of each of these conditions revealed no significant clusters relative to zero.

## Results

### Behavior

When limiting our high-probability trials to those on the horizontal midline, we observed the same high-probability location speedup as previously reported in Duncan, van Moorselaar and Theeuwes (2023), (*t*(21) = 3.834, *p* < 0.001, *d* = 0.82; Figures 1B & C). As is normal in statistical learning paradigms, we also examined whether intertrial effects were present, and as usual a strong effect of target location repetitions was found (*t*(21) = 4.056, *p* < 0.001, *d* = 0.87). To ensure our statistical learning effect was not entirely driven by intertrial priming, the main analysis was also done while excluding all repetition trials (~14% of total trials). This analysis yielded the same pattern of results (*t*(21) = 3.456, *p* = 0.002, *d* = 0.74; note Figure 1B & C do not include repetition trials). Furthermore, nine participants indicated they had noticed at least some level of target regularity. An additional analysis excluding these aware participants showed the speedup effect was not affected by these exclusions (*t*(12) = 2.796, *p* = 0.016, *d* = 0.776). The observation that a clear probability cuing effect was present in our data was sufficient to continue with our neural analysis; for further behavioral analyses on this dataset, including an analysis of distractor trials and neutral trials; see Duncan et al. (2023).

### Lateralized Alpha Results

Separate analyses on lateralized power were done when isolating ping and no-ping trials. Cluster-based analysis of the entire frequency range found no significant clusters in the broadband signal for either trial type when contrasting contra- and ipsilateral electrodes (Figure 2A & B). Furthermore, this analysis when combining the two conditions revealed no significant clusters (Figure 2C). A more targeted analysis of the alpha range in this combined condition similarly found no clusters of interest (Figure 2D, black line. Significant clusters would have been indicated if present). Alpha is occasionally separated between high frequency and low frequency bands (Klimesch, 2012), in fact Wang et al. (2019), found that alpha lateralization in response to statistical learning was primarily present in the lower alpha band (7-10Hz). To rule out the possibility that our alpha band selection was obscuring real lateralization, average alpha in the lower (7-10) and higher (10-13) frequency ranges were also tested for significance. Again, no significant clusters were found in either half of the alpha range (Figure 2D). As a result, the current data provides no evidence that proactive alpha lateralization is a correlate of statistically learned spatial enhancement in the probability cuing paradigm.

### N2pc Results

As visualized in Figure 3, targets presented at the high probability locations appeared to be accompanied by a larger N2pc at high relative to low probability locations. To test these effects, as a first step, N2pc components were measured as the mean amplitudes from 240 – 340-ms post stimulus and subsequently analyzed using a repeated measures ANOVA with within subjects’ factors Target Condition (high-probability, low probability) and Hemifield (contralateral to target, ipsilateral to target). This analysis confirmed that the N2pc was reliable across conditions (main effect Hemifield: *F* (1,21) = 12.217, *p* = 0.002, *η^2^* = .068). Pairwise comparisons between contralateral and ipsilateral waveforms yielded reliable differences in both high- and low-probability target conditions (*t*(21) = 5.586, *p* < 0.001, *d* = 1.191; *t*(21) = 1.842,*p* = 0.04, *d* = 0.393 respectively, one-tailed t-tests). However, the apparent difference between targets at high and low probability locations was not reflected in a significant interaction (*F* (1,21) = 2.539, *p* = 0.126, *η^2^* = .004). As visual inspection showed that the effect was primarily located in the early half of the identified N2pc window, we repeated the analysis separately for the early and the late part of the N2pc component (240-290ms and 290-340ms post stimulus onset respectively). These analyses again yielded reliable N2pc’s across conditions (main effect Hemifield: all *F*’s > 5.4, all *p*’s < 0.03), but critically a significant interaction reflecting a more pronounced N2pc at high vs. low-probability locations was only observed in the early time window (*F* (1,21) = 5.808, *p* = 0.025, *η^2^* = .009).

**Figure 3:**
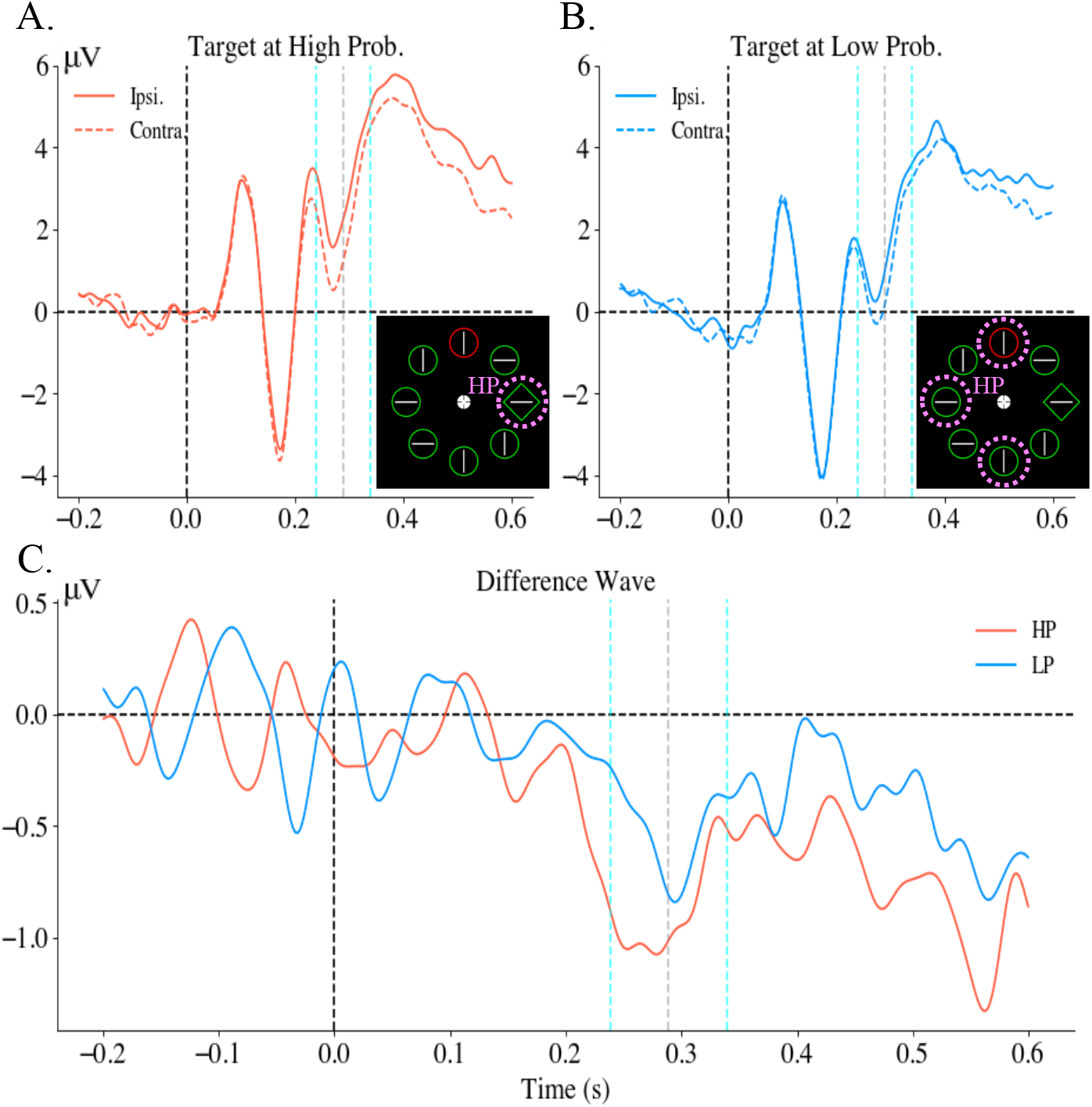
Grand averaged ERP’s of electrodes of interest following search onset (vertical dotted black line) for high-probability (HP) (A) and low probability (LP) (B) trials. In both figures, the solid line represents ipsilateral electrode activity, while the dotted lines represent contralateral activity. Trials pooled to generate these ERP’s always held the target on the horizontal midline. Distractors could be present on the vertical meridian, or were absent. C) Difference waves calculated by subtracting contralateral from ipsilateral evoked activity for HP (red) and LP (blue) trials. Light blue dotted lines in each figure demarcate the N2pc window of interest used in the analysis. Shaded area represent standard error of means. In all figures, the turquoise vertical lines represent the N2pc window identified by the combined peak N2pc amplitude. Grey vertical line represents the peak amplitude point, with the area to the left of the line being the early-N2pc window, and to the right being the late-N2pc window.

## Discussion

Several studies have shown that following statistical learning, visual search is optimally adapted to the spatial probabilities underlying target presentation in a search task (Geng & Behrmann, 2002, 2005; Huang et al., 2022; Zhang et al., 2022). The present study investigated the electrophysiological correlates of learned attentional enhancement, examining how probability cuing modulated two well studied attentional encephalographic indexes: alpha lateralization and the target evoked N2pc ERP component. To this end, we modified the additional singleton paradigm (Theeuwes, 1991, 1992) such that the target appeared with higher probability at periodically shifting locations in the search display. The behavioral data showed more efficient attentional selection of the target when presented at high probability target locations relative to low probability locations confirming earlier findings using this paradigm (Gao & Theeuwes, 2020a; Huang et al., 2022; Zhang et al., 2022) and demonstrating that this learning even occurs when high-probability locations periodically shift during an experimental session. Critically however, this learned attentional enhancement was not characterized by known encephalographic markers of top-down attentional selection – lateralized anticipatory alpha relative to the expected target location. Instead, an exaggerated N2pc at high-probability target locations relative to low probability locations was found, resembling results typical of studies which systematically varied target salience relative to background elements - or bottom-up attentional capture (Berggren & Eimer, 2020; Mazza et al., 2009; Töllner et al., 2011). Such a depiction helps to lay the groundwork for a unified account of selection history effects; one in which history effects are primarily enacted via increased saliency on the attentional priority map.

The N2pc is a well-studied ERP component known to index attentional processing of lateral stimuli (Eimer, 1996; Luck & Hillyard, 1994). In the current study, an exaggerated N2pc was found following targets presented in the high-probability location, localized in in the early half of the N2pc window identified via average peak latency (see methods). This early window stretched from 240-290ms post stimuli onset, falling squarely within what may be considered the standard N2pc window of 200-300ms (Luck, 2014). Furthermore, this early window entirely encompassed the P2 and N2 modulation, a modulation which past studies found to drive the N2pc (Eimer, 1996; Hickey et al., 2006, 2009; Luck & Hillyard, 1994). As a result, we consider this early-phase N2pc to be roughly comparable to what others have called the N2pc in the past.

While traditionally it was believed that the N2pc reflected attentional selection (Eimer, 1996; Woodman & Luck, 1999), more recently it was proposed that this component may in fact track target individuation from background distractors (Mazza et al., 2009, 2013; Mazza & Caramazza, 2015; Pagano & Mazza, 2012). Importantly, the amplitude of the N2pc component has not been shown to modulate as a result of top-down covert orienting (Foster et al., 2020; Kiss et al., 2008). By increasing the relative salience of targets versus nearby competing distractor elements, on the other hand, leads to a monotonic increase in the N2pc amplitude (Berggren & Eimer, 2020; Mazza et al., 2009; Töllner et al., 2011; Zhao et al., 2011), reflecting in part the ease with which target saliency contrasts allows the individuation of the target from background elements. Such an interpretation would suggest that the heightened N2pc observed in the current study reflects an upweighting of saliency information at the learned high-probability target location, resulting in more salient targets evoking greater pop-out effects during parallel search.

It is also well established that the topography of alpha power becomes tuned to expected target locations in response to explicit cues already before search display onset (Foster et al., 2017; Sauseng et al., 2005; Foster & Awh, 2019), While there is some debate on the exact neural mechanism behind this preparatory shift in alpha (Jensen, 2023; Jensen & Mazaheri, 2010), its ubiquity as a marker of covert attention is unassailable. Given the behavioral evidence that statistically learned target enhancement is implemented proactively, for example from capture-probe paradigms (Huang et al., 2021, 2022), there was reason to assume that learned attentional enhancement should also be reflected in preparatory alpha power. At the same time, it should be noted that to date there is actually little support that alpha-band activity is modulated by spatial imbalances across displays (Ferrante et al., 2023; Moorselaar & Slagter, 2019; van Moorselaar et al., 2021; but see Wang et al., 2019). These studies however all examined alpha modulations in response to learning about distractor probabilities, leaving open the possibility that learning about probable target locations is reflected in lateralized alpha band activity before search display onset. Indeed, growing evidence suggests that alpha mediates direct enhancement of input at attended locations, rather than that it suppresses irrelevant input as traditionally assumed (Foster & Awh, 2019; Jensen, 2023). However, the current results did not show reliable tuning within the alpha-band towards in this case the high-probability target location, strongly suggesting that learned attentional tuning is subserved by a different neural mechanism than top-down attentional orienting induced by endogenous cueing.

The current results agree nicely with those recently reported by Ferrante et al. (2023) using a distractor suppression paradigm. In their results, it was found pre-stimulus alpha did not index the learned suppression effect - thus agreeing with several other recent papers (Moorselaar & Slagter, 2019; Noonan et al., 2016; van Moorselaar et al., 2020, 2021) - but statistical learning did lead to reduced proactive excitability of visual neurons in the suppressed hemifield. Ferrante et al. then argue that this reduced excitability may be evidence for a latent synaptic mechanism, suppressing excitability of spatially tuned neurons in suppressed regions. Such a reduction in excitability of spatially tuned neurons may then lead to attenuated saliency signals from the suppressed region in space, leading to the reduced distractor interference. This down-weighting of space on the latent spatial priority map would then represent the opposite effect as what has been reported here, an upweighting of saliency signals for targets presented at enhanced locations in space. It is important to note that just as in the Ferrante et al. (2023) study, the current data are also in line with the idea that the priority landscape, although seemingly only apparent after search display onset, was already in place in anticipation of search display onset. As reported in Duncan et al. (2022) multivariate analyses yoked to the neutral pings presented on half of the trials before search display onset revealed the anticipatory priority landscape. It thus appears that that while statistical learning proactively adjusts the spatial priority map, this priority landscape only becomes apparent after integration of bottom-up sensory input (van Moorselaar & Slagter, 2020) such as a probe display (Huang et al., 2021, 2022), a task irrelevant perturbation (Duncan et al., 2022; Ferrante et al., 2018) or as reported here the onset of the actual search display.

Intriguingly, an exaggerated N2pc such as that seen in the current data is not just known to present itself in situations of heightened target-background salience (Mazza et al., 2009; Töllner et al., 2011), but also in the context of contextual cuing paradigms (Johnson et al., 2007; Olson et al., 2001; Schankin & Schubö, 2010; Zinchenko et al., 2020) and value driven attentional capture paradigms (Hinault et al., 2019; Kiss et al., 2009; MacLean & Giesbrecht, 2015; Qi et al., 2013). Thus, the ubiquity of exaggerated N2pc’s in the context of selection history effects suggests this neural marker may be a general correlate of selection history driven attentional enhancement. However, while probability cuing is known to be at least partially proactive in nature (Duncan et al., 2022; Ferrante et al., 2023; Huang et al., 2021, 2022), contextual cuing and value driven capture are necessarily reactive, and any unified neural theory of selection history would struggle to reconcile these differences. One possibility is that these numerous effects are mediated by priority maps on various feature dimensions (Found & Müller, 1996; Müller et al., 1995; Zelinsky & Bisley, 2015). The facilitation or suppressed of feature tuned neurons via changes in neural excitability would then lead to changes to a stimulus’ perceived salience, thereby influencing behavior. Clearly much more work is needed to continue illuminating how selection history influences perception to guide future attentional deployment.

Finally, both the finding that common electrophysiological markers of top-down attention are absent in EEG recordings of a probability cuing paradigm, as well as the finding that similar N2pc characteristics arise among various known selection history effects motivates the adoption of the new tripartite model of attention first proposed by Awh and colleagues (2012). While some have instead suggested a subsuming of selection history and top-down effects into a single category of the sake of convenience (Egeth, 2018; Gaspelin & Luck, 2018), the current study adds to the growing evidence that selection history in fact represents a distinct cognitive mechanism from top-down attention – localized in different brain areas (Geng & Behrmann, 2002; Schapiro et al., 2014; Shaqiri & Anderson, 2012, 2013), exhibiting different behavioral tendencies (Gao & Theeuwes, 2020a, 2020b; Geng & Behrmann, 2005; Jiang et al., 2013; Wang & Theeuwes, 2018; Won & Jiang, 2015), and now having been shown to exhibit distinct electrophysiological markers consistent with distinct neural mechanisms underlying the attentional effects. In fact, if anything, the current results suggest that selection history effects may be thought of as essentially more ‘bottom-up’ in nature than ‘top-down’ (see Theeuwes, 2018 for a similar argument).

## Acknowledgments

This research was supported by a European Research Council (ERC) advanced grant 833029 – [LEARNATTEND].

## Footnotes

1 The two excluded participants had HP presentation orders of [Left, Bottom, Right, Top] and [Right, Bottom, Left, Top].

